# ProteinDock: A physics-informed layer to improve protein-protein docking reliability

**DOI:** 10.64898/2026.07.17.739238

**Authors:** Gowrish Rajagopal, Søren C. Spina, Joseph S. Bailey, Blaise R. Kimmel

**Author notes:** To whom correspondence should be addressed **Corresponding Author** Blaise R. Kimmel, Ph.D. 151 W. Woodruff Avenue Columbus, OH 43210.

## Abstract

Computational modeling provides geometric insight into protein-protein interactions without requiring the resources of experimentation. However, reliability can be hindered when modeling proteins with distinctive features, such as antibodies, that use flexible, polar-rich loops to bind antigens. We developed *ProteinDock*, a physics-based tool that can be used in combination with leading modeling programs to improve the reliability of protein-protein docking; this work provides a case study of antibody-antigen interfaces. *ProteinDock* was layered onto Rosetta for docking unbound experimentally determined structures, and when evaluated on Docking Benchmark Set 5.5, generated CAPRI acceptable-quality or better for 80.2% of targets, an improvement of 32.8 percentage points over vanilla Rosetta’s 47.4% on the same dataset. To improve protein-protein prediction reliability from sequence inputs, we demonstrate that a truncated version of ProteinDock can be used to choose the optimal prediction among outputs from multiple deep learning-based tools. We show that this strategy is a computationally efficient alternative to increasing the seed quantity for deep-learning predictions. A graphical user interface for layering *ProteinDock* has been created and is available at https://github.com/Kimmel-Lab/proteindock and https://proteindock.com/.

**TOC Figure:** 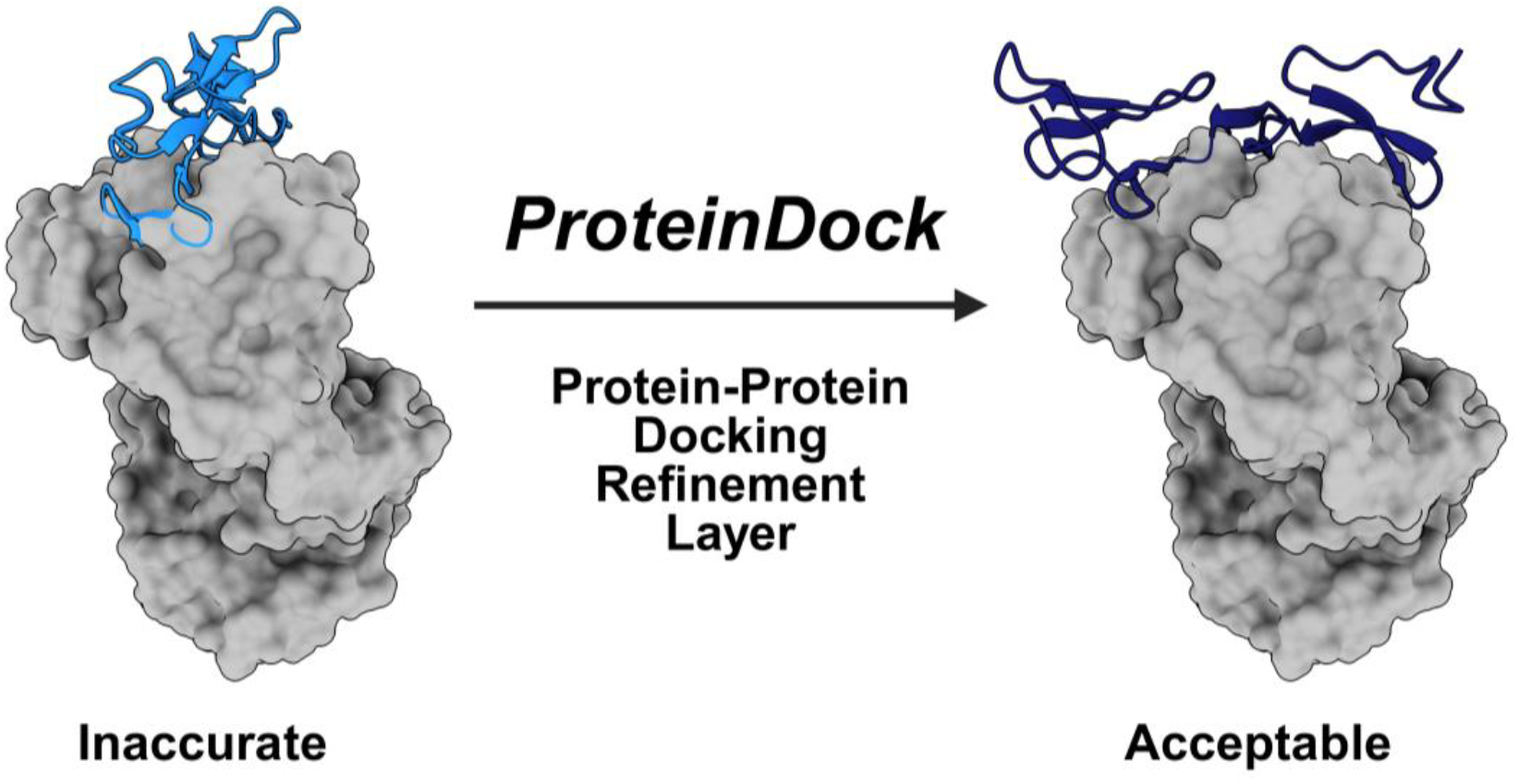

## Introduction

Proteins interact with one another to drive essential biological processes such as immune function, muscle contraction, and cell division. The interface chemistry of large, multi-protein complexes makes these interactions geometrically specific, and three-dimensional (3-D) structural models are a necessary tool for observing, learning from, and designing protein-protein interactions, particularly for engineering new binders and bioconjugates^1,2^. For over half a century, experimental determination has been the primary method for obtaining “ground-truth” structural models of protein-protein complexes^3^. However, experimental methods are costly and time-consuming, motivating the use of computational tools to predict protein-protein complexes^4^.

Computationally docking two experimentally determined structures can obviate the need to experimentally determine a bound complex. A leading tool for this task, Rosetta, uses the ref2015 energy score to dock complexes based on physics-based principles, and this strategy is largely effective for modeling general protein-protein binding^5^. Modeling interaction types with unusual characteristics can reduce reliability, motivating the use of modified score functions optimized for specific interaction types, such as protein-peptide binding^6^. With Rosetta used to dock two structural model inputs, the use of these modified score functions can be constrained to proteins that have been experimentally determined previously.

AlphaFold3 (AF3) allows users to predict singular proteins and protein complexes from sequence inputs alone, circumventing the need for experimentally resolved structures^7^. This allows modeling of unsolved proteins, engineered proteins, and proteins for which all experimentally sourced structures are in bound states, capabilities that were previously impossible. However, structural models predicted from sequence inputs alone consistently perform less reliably than those with resolved experimental structures due to inaccuracies such as minor differences in sidechain packing. While increasing the seed count is a strategy shown to improve AlphaFold3 reliability, computational cost increases significantly with each additional seed. *AlphaRED* (AlphaFold-initiated Replica Exchange Docking) was developed by fusing the physics-based ReplicaDock2.0 with the deep-learning-based AlphaFold-multimer to reduce incorrect AlphaFold complex predictions at minimal computational cost^9^.

Motivated in part by *AlphaRED’s* success in adding a physics-based layer to improve the prediction reliability of AlphaFold-multimer, we have developed a physics-based layer, *ProteinDock*, to improve the protein-protein modeling reliability of existing tools, evaluated on antibody-antigen test sets. *ProteinDock* Mode 1 was developed to improve the reliability of protein-protein docking when layered onto physics-based tools that use unbound, experimentally resolved structures as inputs. *ProteinDock* Mode 2 was created to improve the reliability of protein-protein complex predictions by deep-learning models given only sequence inputs. Evaluated on antibody-antigen test sets, *ProteinDock* “improves reliability” when eliminating docking “failures”, using the fraction of targets reaching acceptable or better quality per DockQ scoring as the guiding metric. For ease of use and widespread accessibility, a graphical user interface has been created for layering *ProteinDock* and is accessible at https://github.com/Kimmel-lab/proteindock and https://www.proteindock.com.

## 2. Methods

### Benchmarks

We evaluated *ProteinDock* on three benchmarks: Docking Benchmark version 5.5 (DB5.5)^10^, Structural Antibody Database-Expanded (SAbDab-EX; branched from original SAbDab database)^11^, and novel-50^12^. DB5.5 is a standard unbound protein-protein docking dataset, SAbDab-EX provides unbound antibody-antigen docking cases, and the novel-50 set consists of 50 antibody-antigen complexes that were entered into the Protein Database after AlphaFold3’s training cutoff date. Full dataset filtering details for all three datasets, resolution thresholds, structural relaxation cutoffs, and date ranges are given in **Supplementary Table 1**. Unlike DB5.5 and SAbDab-EX, novel-50 is not used for direct method comparison in Mode 1. It tests the ability of *ProteinDock* to generalize its refinement to post-cutoff antibody-antigen targets.

To encourage generalization and control inflated performance caused by memorization, protein targets are typically filtered at < 30% sequence identity^13^. This is not achievable with antibody benchmarks due to the immunoglobulin framework shared across antibodies^14^. All novel-50 candidates have more than 30% identity with each other (median 84%), reflecting this shared biology.

Because docking predicts the binding between two chains and does not directly involve the structure of either chain alone, we define **pair-novelty**, the absence of a specific antibody-antigen pair in the PDB prior to the AlphaFold3 cutoff, as the main novelty measure throughout this work (computation described in Supplementary Table 2). 42 of 50 candidate pairs in novel-50 are found to be pair-novel. The other 8 share sequence similarity with pre-cutoff antibodies are post-cutoff SARS-CoV-2 spike-antibody complexes. We chose to keep all eight because their sequence similarity to pre-cutoff antibodies does not imply the specific pairings were seen in training, and their performance is reported separately in Results.

*ProteinDock* is implemented in Python 3.10 with *PyRosetta4*, using the ref2015 scoring function^13– 15^. The electrostatic term (fa_elec) is increased by 1.5x to account for the electrostatically-driven nature of antibody-antigen interfaces, with all other ref2015 terms unchanged^16^. We assessed the sensitivity of *ProteinDock’s* performance to the value of the fa_elec’s reweight value for both Mode 1 and Mode 2 (Results).

*ProteinDock* has two modes. Mode 1 does local rigid-body docking with all-atom interface refinement on two chain coordinate inputs (Supplementary Table 2). It takes unbound monomer structures from DB5.5 and SAbDab-EX or bound-conformation chains extracted from complexes for novel-50 (coordinate-given inputs). Mode 2 does not perform sampling but arbitrates between top-1 poses of two independent sequence-based foundation models (AlphaFold3 and Boltz-2), using *Rosetta*’s ref2015-based interface energy to choose between them (**Figure 1**)^17^.

**Figure 1.**
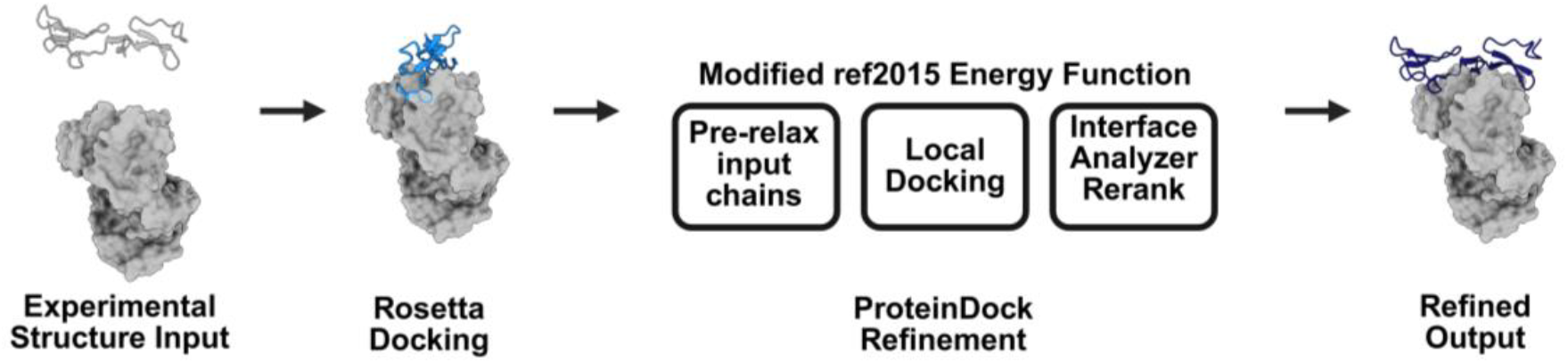
Workflow schematic of *ProteinDock* Mode 1 improving the reliability of protein-protein docking with Rosetta.

### Implementation

#### Mode 1 (Layer for Docking Tools)

Unbound structures for *DB5*.*5* and *SAbDab*, and bound-frame chains for novel-50 are the two independent monomer coordinates used as inputs for the pipeline. Each chain is first independently pre-relaxed with *FastRelax* (1 cycle) using the ref2015 score function, then 50 structures are docked locally using Rosetta’s *DockingProtocol* under the increased electrostatic term weight (fa_elec x 1.5)^18^. The predicted pose with the top Rosetta interface score (I_sc) is taken from the docking scorefile. The single relax cycle is due to optimizing for speed, chosen to keep per-decoy relax feasible at nstruct=50^18^. Results confirm that this does not limit performance.

For direct comparison, *DB5*.*5* and *SAbDab* inputs are the same unbound monomer coordinates across all methods (*ProteinDock*, HADDOCK, and baseline Rosetta). The novel-50 dataset has differing inputs for both *ProteinDock* and tools like *AF3* and *Boltz-2. ProteinDock’s* Mode 1 receives bound-frame chains taken directly from the complex, while AlphaFold3 and Boltz-2 receive sequences only. We made this choice for two reasons. First, unbound monomer structures do not exist for post-cutoff targets, as only the bound complex has been solved for more recent structures (such as the ones found in the dataset). Secondly, bound-frame chains match the inputs seen in real antibody-design workflows^20^, where designed antibodies and antigens are already available in a bound-like conformation, only needing rigid-body placement and interface refinement to finish the process.

Mode 1 novel-50 results are not intended to be benchmarked against sequence-based foundation models; they are intended to analyze *ProteinDock’s* placement and refinement of two bound-like chains into their native complex. *ProteinDock’s* operating range is further detailed in the capture-radius analysis (**Supplementary Methods 1 and Supplementary Table 3**). To check whether bound-frame inputs are inflating results, we separately ran the same process on unbound inputs for a subset of novel-50 targets (Supplementary Table 3). The direct comparison to the sequence-only predictors is made in Mode 2, where *ProteinDock* scores the same predicted coordinates as AlphaFold3.

#### Mode 2 (Layer for Structure Prediction Tools)

Mode 2 reframed *ProteinDock* as a scoring layer for outputs of foundation models, after finding that direct redocking or refinement of foundation model poses failed on improving original outputs. The candidate poses came from *ProteinDock* layered onto either AF3 (PD-AF3) or a combination of AlphaFold3’s and Boltz-2’s top-1 structures (AF3-Boltz-PDT). In both *ProteinDock* layered variants, candidates were scored with PyRosetta’s *InterfaceAnalyzerMover* (Chaudhary *et al*., 2010) under ref2015 with *fa_elec* x 1.5. The function was configured for scoring only (pack_separated=False, pack_input=False, compute_interface_sc= False, compute_packstat= False) and final selection was based on argmin dG_interface (**Table 1**), (ΔG) the Rosetta calculated interface binding energy. Because repacking would move the predictor’s own side chains, no repacking, *FastRelax*, or rigid-body sampling was run prior to scoring, avoiding any changes *DockQ* could no longer attribute to the predictor itself. Mode 2 inputs were foundation-model predicted poses (AlphaFold3 and Boltz-2). They did not include any confidence metrics, MSAs, or template signals, and did not apply any per-target hyperparameters or confidence thresholds (**Figure 2**).

**Table 1.** Mode 1 and Mode 2 Variants used in this work.

| Variant | Mode | Input | Sampling | Selection | Role |
| --- | --- | --- | --- | --- | --- |
| PD | 1 | Unbound or bound-frame monomers | Pre-relax+ local docking (nstruct=50) @ fa_elec x 1.5 | Top-1 by initial l_sc | *Baseline ProteinDock* docking method |
| PD-AF3 | 2 | AF3's 25 seed x sample poses | None (Scoring only) | argmin dG_interface | Single-predictor rescoring |
| AF3-Boltz-PDT | 2 | AF3's top-1 + Boltz-2's top-1 | None (Scoring only) | argmin dG_interface with fallbacks | Cross-predictor arbitration |

**Figure 2.**
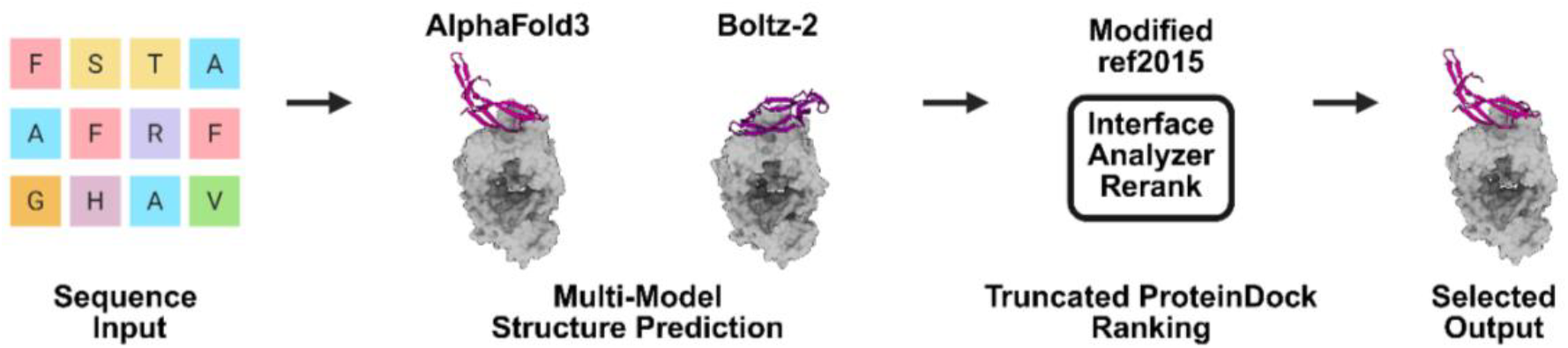
Workflow schematic of truncated *ProteinDock* Mode 2 improving the reliability of complex structural prediction by reranking outputs of multiple deep-learning tools.

PD-AF3 and the AF3-alone baseline used the same 25 predicted structures per target and differed only in the selection methods. AF3 structures were default ranked by argmax ranking_score (AlphaFold3’s internal confidence-based rank for structures), while PD-AF3 decided via argmin dG_interface. AF3-Boltz-PDT applied a more conservative selection rule, because it never lets dG_interface override AlphaFold3 outright, and only abandons it fully if no output was produced. We computed dG_interface for AF3’s top ranking structure and the Boltz predicted complex with the highest ‘confidence score’ (a composite score of pLDDT and predicted interface accuracy), and chose whichever gave the more favorable value. If Rosetta was unable to converge, and thus unable to score an AF3 predicted structure, we kept the AF3 pose regardless, since Rosetta scoring failures limited *ProteinDock* inputs for Mode 2 rather than providing concrete evidence against the predictor. If AlphaFold3 was unable to produce output at all (an inference crash), we fell back to the top Boltz-2 prediction. There were seven other Mode 1 variants (PD+, PD-AF3-Split, PD-AF3-Split+, PD-AF3-Polish, PD-AF3-Refine, PD-AF3-Cluster, PD-AF3-Dock) that were evaluated as potential ablations. Their results and the reasons they are not being highlighted are discussed below and fully specified in **Supplementary Table 4**.

### Components

*ProteinDock* Mode 1 protocol adds two additional steps to the baseline Rosetta docking architecture, including a pre-relax step, and the fa_elec x 1.5 reweighting. To understand the contribution of each, we test two additional conditions, one with only the reweighting applied and the other with only the pre-relax step (**Table 2**).

**Table 2.**
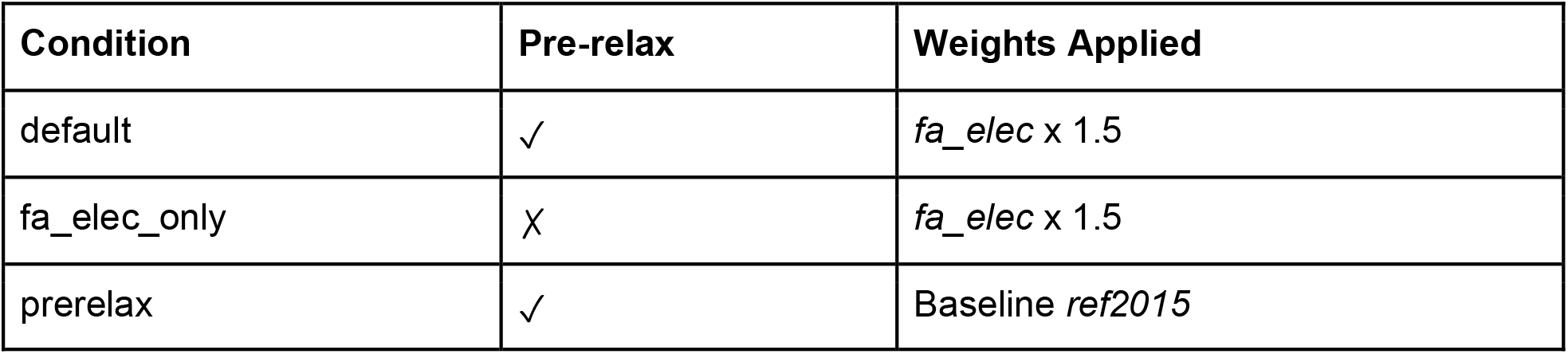
Component ablation conditions and weights applied in this study.

### The capacity ceiling of ref2015-feature scoring

We tried boosting the scoring signal for ref2015 to assess its ceiling using techniques such as physics-based weight corrections and deep learning models (**Supplementary Methods 2**). We swept the model capacity across roughly four orders of magnitude over a pool of 13,240 DockQ-scored decoys (**Supplementary Table 3**), under 5-fold PDB-disjoint cross-validation. This was done across four model classes: 14 single-parameter corrections; a 7-parameter linear refit of ref2015 energy terms under pairwise margin-rank loss; 27 conventional regressors (Ridge, gradient-boosted, random forest) over 27 interface geometry features; and a 15-seed set-transformer ensemble. Full architecture and training details are in **Supplementary Table 4**.

### Scoring and Metrics

The predictions from all the Mode 1 and 2 variants were scored with the DockQ v2 Python package^21^. We classified DockQ scores into the four CAPRI quality bins: High (≥ 0.80), Medium (0.49-0.80), Acceptable (0.23-0.49), and Incorrect (< 0.23)^22^. The main metric reported throughout the work is success rate, the proportion of targets reaching the acceptable bin or higher (DockQ ≥ 0.23). The oracle (a selector that always picks the highest-DockQ decoy for each target, regardless of any score the pipeline itself computes) selected the highest-DockQ decoy for each target to establish the selection ceiling, which is reported for Mode 1 and for AF3’s 25-sample pool. When a target has no residues within 10 Å of any native interface residue, DockQ returns iRMS = 999. These iRMS=999 scored targets were counted as Incorrect (DockQ < 0.23) in the pipeline for success rate and are differentiated from missing-output failures (when a method produced no scorable structure). In the reliability-layer evaluation of Mode 2, candidate poses that received no dG_interface score due to a missing chain remained in the denominator of the output and were recognized as method failures. AlphaFold3-alone, Boltz-2, PD-AF3, and AF3-Boltz-PDT were evaluated under the same DockQ pipeline on the sequence-only novel-50 set. Native structure preparation and chain-merge corrections are described in **Supplementary Table 2**.

### Statistical Tests

Confidence intervals for single success-rate values were computed using both Wilson 95% binomial intervals and 10,000-resample target-level bootstrap intervals, because bootstrap resampling can fail with small sample sizes. Wilson is used throughout the main text.

### Baselines

*ProteinDock* was evaluated against baseline packages in both structure-folding and docking categories. AlphaFold3 and Boltz-2 were the sequence-based folding baselines, and HADDOCK3 and standalone Rosetta were the physics-based docking baselines. We ran the public AlphaFold3 release in protein-only mode^7^. The sample with the highest ‘ranking_score’ was taken as the per-target prediction. On DB5.5 and SAbDab-EX, inference used a single random seed with 5 diffusion samples and on novel-50, we ran 5 independent seeds with 5 diffusion samples each. The AlphaFold3 template database was filtered to match the model’s training cutoff. We verified that the predicted binding geometry was absent from both the training and template databases, even when a pre-cutoff template was retrieved for an individual chain. We used Boltz-2 v2.2.1 with one random seed per target and MSA (multiple sequence alignment) construction ran through the ColabFold MMseqs2 server^23^, where we took the top ranked sample as the per-target prediction.

Due to the availability of interface information, we ran HADDOCK3 under different variants of the program that differ in availability of interface information^23^. To differentiate the performance between HADDOCK3 and *ProteinDock*, two main variants were run. These variants reflect improvements due to interface information versus the docking algorithm itself, using same chain coordinates, mimicking local *ProteinDock* inputs, and using native residues as ambiguous interaction restraints (AIR). We ran HADDOCK3 v2025.5.0 on DB5.5 with each variant and selected the top structure by HADDOCK score. We ran baseline Rosetta local docking parameters on DB5.5, using the same input style as the *ProteinDock* Mode 1 protocol to understand the contributions of the different components of *ProteinDock*. Unlike Mode 1 which featured a pre-relax step, this baseline was run with no pre-relax step and scored with default ref2015 weights to help understand the specific contribution of each components within *ProteinDock*. Full inference parameters, hardware, and runtime details are given in **Supplementary Table 5**. All Mode 1 and Mode 2 variants were run on a single Intel Xeon Gold 6148 core (4 cores per SLURM array task) on the Pitzer/Cardinal clusters of the Ohio Supercomputer Center. The foundation model predictors of Mode 2 (AlphaFold3, Boltz-2) were run on NVIDIA A100 GPUs in the same cluster, while the arbitration step was run on the aforementioned CPU core.

## Results

ProteinDock Mode 1 is a refinement layer that improves the reliability of protein-protein modeling in physics-based docking tools that use structural inputs (**Figure 3A**). On the Docking Benchmark Set 5.5, which includes various protein and interaction types, Rosetta with the *ProteinDock* layer (Rosetta-PD) generated acceptable-quality or better quality (success rate), for 80.2% of targets, a significant improvement from the 47.4% success rate of standalone Rosetta and 9.1% success rate of HADDOCK (**Figure 3B**). On DB5.5, AlphaFold3’s top-1 success rate drops from 76.2% (186/244) on non-antibody-antigen targets to 53.8% (14/26) on antibody-antigen targets (Fisher’s exact, p = 0.019), showing AlphaFold3 struggles specifically with antibody-antigen pairs. Wilson 95% binomial confidence intervals for every reported variant are given in **Supplementary Figure 1**. As a case study, we evaluated *ProteinDock* on a set of antibody-antigen pairs, SAbDab-EX, where it achieved success for 74.1% of targets, another significant improvement over the 0.0% success rate of HADDOCK, which had the same information available as *ProteinDock*, and a modest improvement over the 65.5% success rate of standalone Rosetta. *ProteinDock* also approaches the success rate of HADDOCK3 given the native interface as ambiguous interaction restraints (78.6%), despite receiving no such inputs. HADDOCK3 performance across its different forms on both benchmarks is shown in **Supplementary Figure 2**. *ProteinDock* Mode 1 requires a computational cost that is twice that of baseline Rosetta (**Figure 3C**).

**Figure 3.**
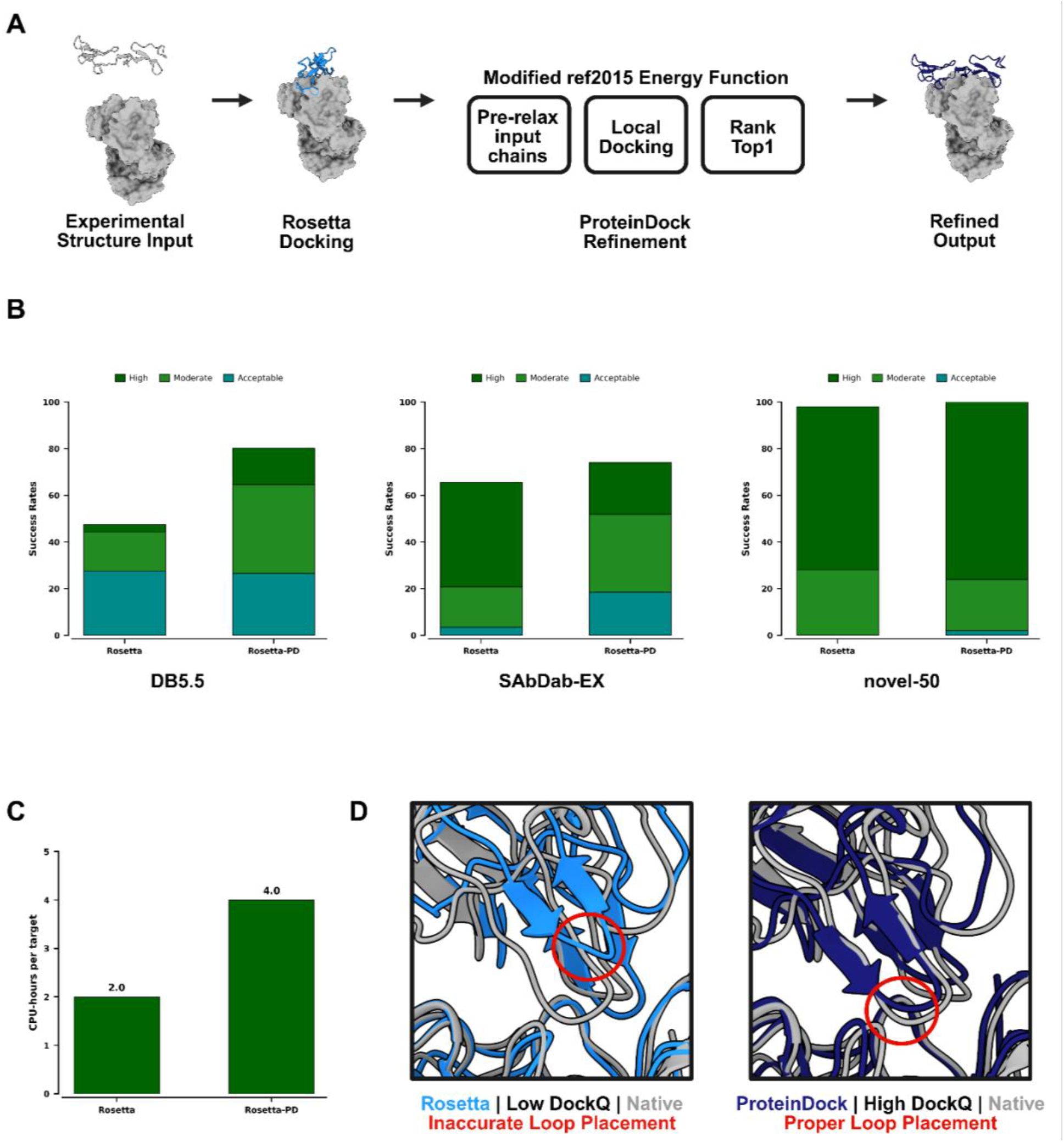
*ProteinDock* Mode 1 can be layered onto physics-based docking tools that use unbound structural inputs.(A) Schematic workflow of truncated *ProteinDock* Mode 1 and use with Rosetta. (B) Evaluation of *ProteinDock* on DB5.5 (n=271), SAbDab-EX (n=29), novel-50 (n=50). Success rates are defined by High (DockQ ≥ 0.80), Moderate (DockQ < 0.80 and ≥ 0.49), Acceptable (DockQ < 0.49 and ≥0.23), Failure (DockQ < 0.23). (C) Computational cost comparison of Rosetta and Rosetta with the *ProteinDock* layer. (D) *ProteinDock*-layered Rosetta (dark blue) had a higher DockQ score and, modeled interface loops with closer resemblance to the experimentally validated native structure (grey), PDB: 1IQD^25^, than baseline Rosetta (light blue).

When experimental structures of unbound proteins are unavailable, deep learning-based methods are commonly used to predict protein-protein complex structures from two input sequences alone. We investigated whether a second mode of *ProteinDock* could be developed to improve the reliability of antibody-antigen predictions by AlphaFold3 (AF3), a deep-learning-based model. The first iteration of *ProteinDock* Mode 2 takes the outputs of deep-learning-based models and processes them through the steps of Mode 1. When this setup was evaluated on novel-50, single-seed AF3 with the *ProteinDock* layer (AF3-PD) was unable to significantly reduce failures compared to standalone single-seed AF3, with both generating CAPRI acceptable-quality or better for 54% of targets. The distribution of per-target ΔDockQ (PD – AF3) on DB5.5 and SAbDab-EX is shown in **Supplementary Figure 3**.

Previous studies have used Rosetta-based binding energy to score antibody-antigen predictions from AF3, and this motivated the next iteration of Mode 2, a truncated version of *ProteinDock* (PDT), strictly consisting of reranking using *InterfaceAnalyzerMover* with the modified ref2015 score function, which chooses the top prediction between AF3 and Boltz-2 outputs (**Figure 4A**). We evaluated this configuration, AF3-Boltz-PDT, on novel-50, where the top-ranked prediction was CAPRI acceptable-quality or better for 60% of targets, a significant improvement over the 54% achieved by standalone single-seed AF3 and the 32% achieved by standalone Boltz-2 (**Figure 4B**). This truncated *ProteinDock* setup is also computationally efficient (**Figure 4C**). These findings show that a physics-based reranking procedure can efficiently evaluate the outputs of multiple tools to determine a top prediction of CAPRI-acceptable quality, with improved reliability compared to a single tool alone (**Figure 4D**).

**Figure 4.**
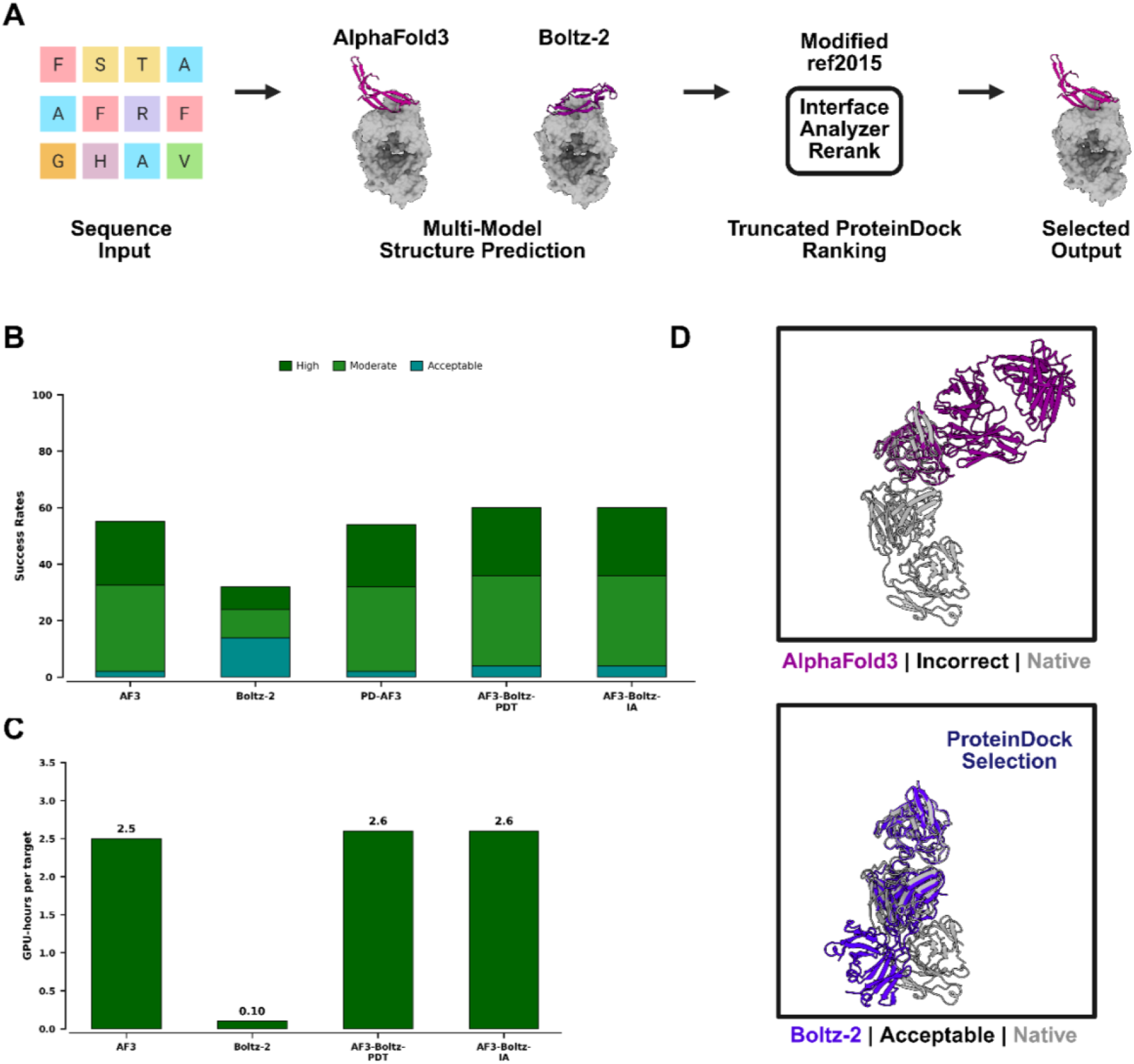
*ProteinDock* Mode 2 can be layered onto deep learning-based tools that predict from sequence inputs alone. (A) Schematic workflow of truncated ProteinDock Mode 2 being used to rank outputs from multiple tools. (B) Evaluation of ProteinDock on novel-50 (n=50). Success rates are defined by High (DockQ ≥ 0.80), Moderate (DockQ < 0.80 and ≥ 0.49), Acceptable (DockQ < 0.49 and ≥0.23), Failure (DockQ < 0.23). (C) Computational cost of workflows. (D) When given structural prediction outputs from AlphaFold3 (magenta) and Boltz-2 (purple), ProteinDock ranked the Boltz-2 model highest, which more closely resembles experimentally validated structure (grey), PDB: 9XYB^26^.

We also evaluated this setup with *InterfaceAnalyzerMover* using an unmodified ref2015 score function (AF3-Boltz-IA), in which the top-ranked prediction was CAPRI acceptable-quality or better for 60% of targets, matching AF3-Boltz-PDT. This comparison indicates that the modification to the ref2015 score function in *ProteinDock* is not responsible for the reliability improvement AF3-Boltz-PDT achieves, however, it does show that a Rosetta-based *InterfaceAnalyzerMover* layer can be used to rank outputs from multiple deep-learning structural prediction models.

### Ablation

We assessed the direct contribution of each component in *ProteinDock* through an ablation test. On a DB5.5 (N=10) subset, we found that pre-relax alone was the major improvement over general blind docking, with the success rate improving from 0% to 90%. The full *ProteinDock* protocol reached 100% success rate, and variants run with fa_elec upweighted beyond 1.5x underperformed. Ablation testing was expanded to the novel-50 dataset. Default Rosetta gave strong results (98% success rate), and both pre-relax alone and fa_elec alone underperfomed the baseline. Only the full combined *ProteinDock* protocol exceeded the baseline results, reaching 100% success rate. The full protocol also improved prediction quality (high-quality bin from 70% to 76%), and adding top-5 reranking additionally improved it further to 82% (**Figure 3**). Per-agreement between PD and PD+ is characterized in **Supplementary Figure 4**. The electrostatic term was pre-registered before the novel-set ablation run. We tested the sensitivity of term across five values between 1.0x and 2.0x for fa_elec on a 20-target DB5.5 subset, which included 5 antibody-antigen complexes and 15 general PPIs. All five reached 100% success rate with fa_elec x 1.5 giving the highest median DockQ score (0.794).

To understand the effects of different designs in the Mode 2 arbitration rescoring layer, we investigated possible designs. Variants that redocked or re-scored AlphaFold3 poses all underperformed the baseline (0-30% success rate), and rescoring AF3’s sample pool without a second predictor yielded similar outcomes. PD-AF3-Split success rate is strongly dependent on the pose quality of AlphaFold’s outputs (**Supplemental Figure 5)**. Only cross-predictor models, reranking both AF3 and Boltz-2 together, improved success rate, reaching 60% with 3 targets being rescued into a successful dock. This did not reach statistical significance at this sample size (McNemar p = 0.25), although the rules of the main arbitration layer guarantee zero loss. Antibody-antibody interfaces were significantly more polar and smaller than non-antibody interfaces (**Table 5**). All five descriptors tested differed significantly between the two groups (p ≤ 0.012).

**Table 3.**
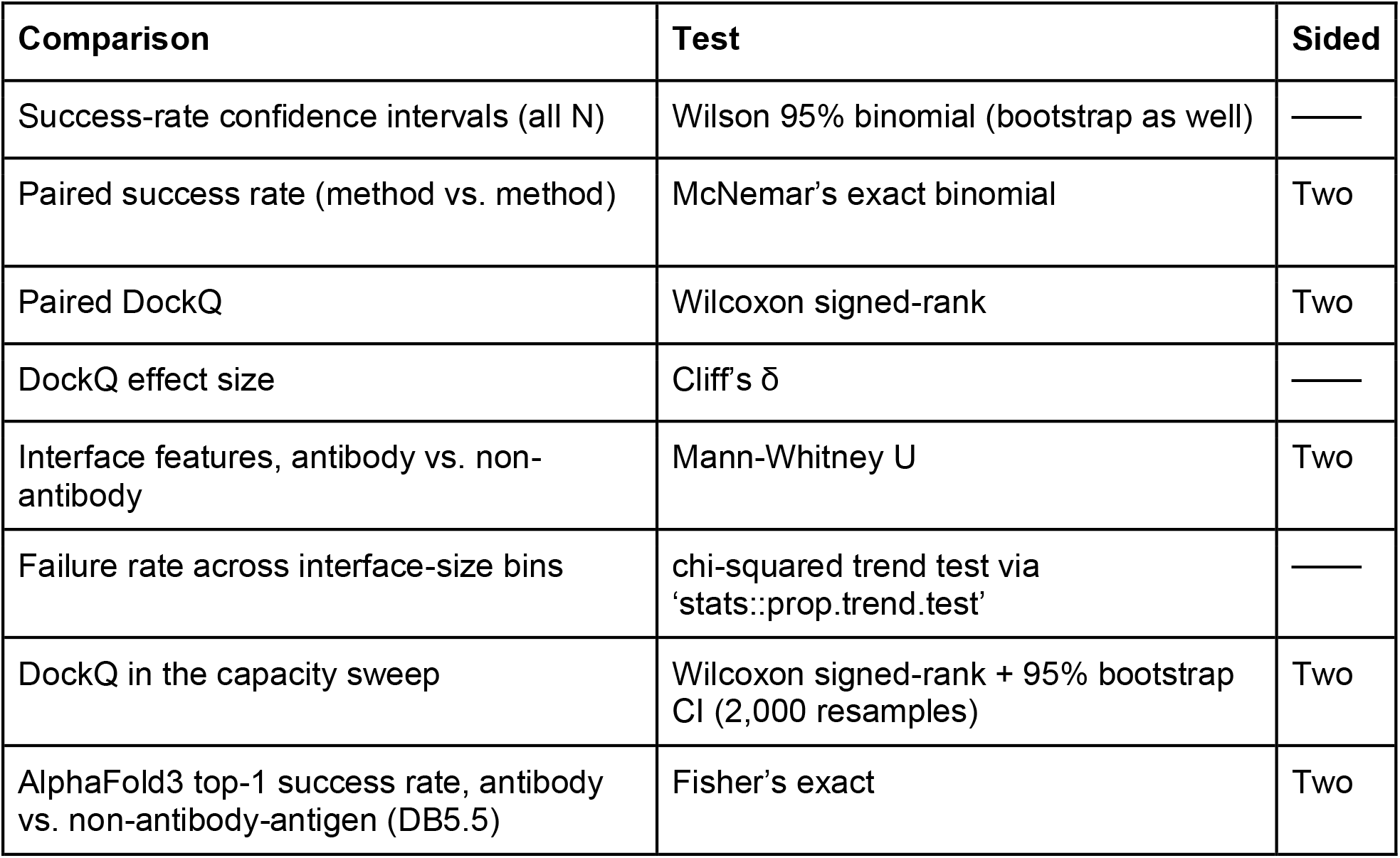
Statistical Tests used in this study.

**Table 4.** Expanded component ablation on novel-50.

| Condition | Pre-relax | fa_elec | Extension | Success Rate (DockQ≥0.23) | Medium+ (≥0.49) | High (≥0.80) | N |
| --- | --- | --- | --- | --- | --- | --- | --- |
| Default Rosetta | — | 1.0× | — | 98.0% (49/50) | 98.0% (49/50) | 70.0% (35/50) | 50 |
| Pre-relax only | ✓ | 1.0× | — | 96.0% (48/50) | 94.0% (47/50) | 78.0% (39/50) | 50 |
| fa_elec × 1.5 only | — | 1.5× | — | 94.0% (47/50) | 90.0% (45/50) | 66.0% (33/50) | 50 |
| Full recipe (PD) | ✓ | 1.5× | — | 100.0% (50/50) | 98.0% (49/50) | 76.0% (38/50) | 50 |
| Top-N rerank (PD+) | ✓ | 1.5× | FastRelax top-5 | 100.0% (50/50) | 100.0% (50/50) | 82.0% (41/50) | 50 |
| AF3-start (PD-AF3-Split)* | ✓ | 1.5× | AF3 input | 38.1% (16/42) | 33.3% (14/42) | 4.8% (2/42) | 42* |

\*PD-AF3-Split N=42: 8 targets excluded due to Rosetta preprocessing failures on AF3 poses; retry did not change outcomes.

**Table 5.** Antibody vs non-antibody interface chemistry.

| Descriptor | Antibody median | Non-antibody median | p |
| --- | --- | --- | --- |
| Hydrophobic fraction | 0.222 | 0.301 | $1.3 \times 10^{-4}$ |
| Polar fraction | 0.391 | 0.320 | $1.1 \times 10^{-5}$ |
| Interface size (residues) | 24 | 32 | $3.1 \times 10^{-5}$ |
| Charged residues | 5 | 7 | $8.1 \times 10^{-8}$ |
| Salt bridges | 0 | 1 | 0.012 |

**Figure 5.**
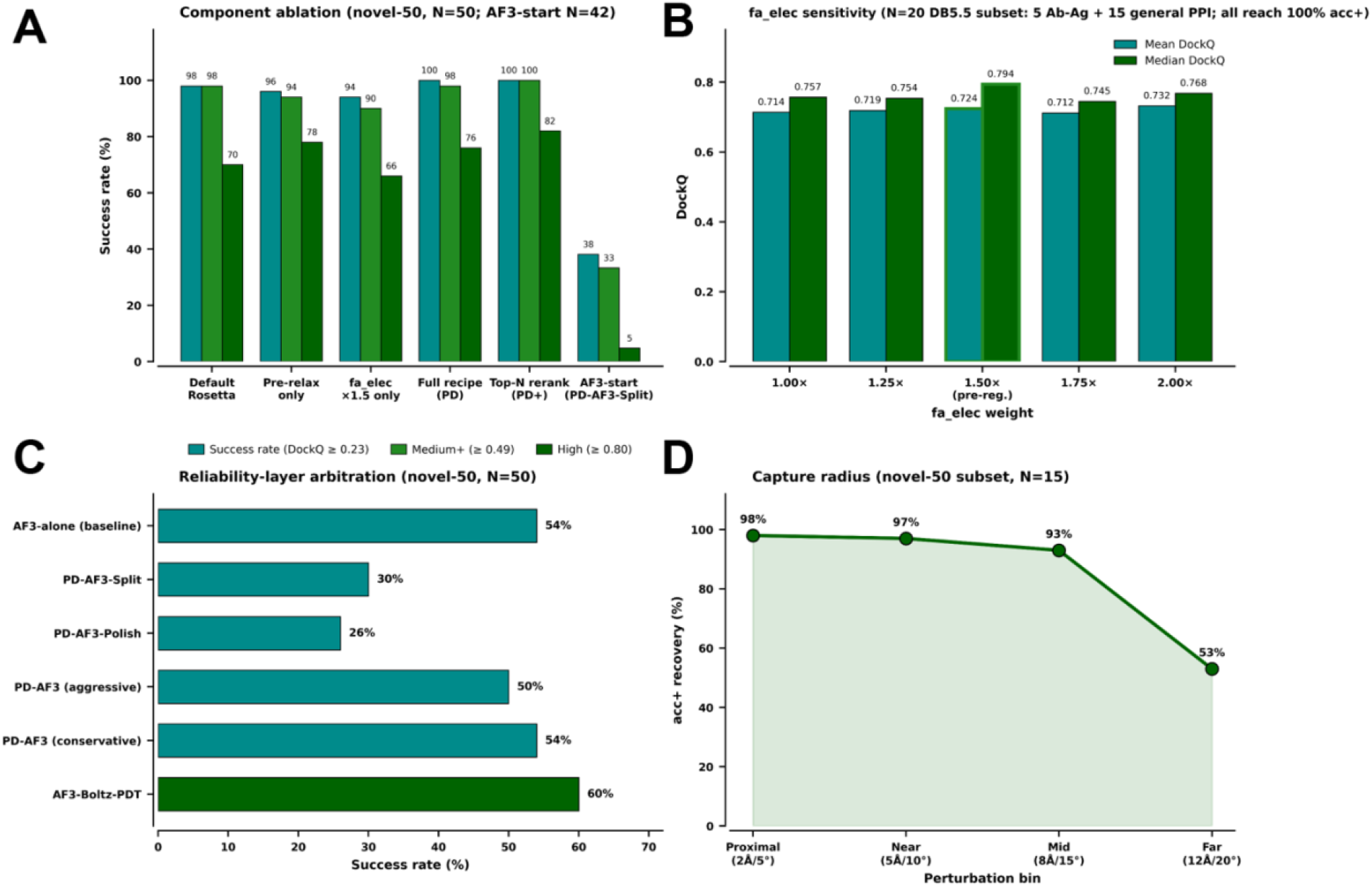
*ProteinDock* mechanism. (A) Component ablation on novel-50, showing success rate at three DockQ thresholds. (B) fa_elec sensitivity sweep on a 20-target DB5.5 vaidation subset across different weights. (C) Reliability-layer arbitration on novel-50 comparing AF-3 alone baseline against different variants. (D) Capture-radius robustness on 15 novel-50 targets across four perturbation magnitudes.

We analyzed how robust the reliability improvement is, given its tolerance to imperfect starting poses and the possibility of training data leakage (**Figure 5**). *ProteinDock* recovered success rate for ≥93% of targets when perturbed up to 8 Å translation/ 15° rotation from the native pose; however, this falls sharply to 53% at 12 Å /20°, effectively forming an 8 Å /15° capture radius. The second check tests whether performance reflects genuine generalization. *ProteinDock* reached 100% success rate on novel-50 across different novelty definitions, arguing against training data leakage as an explanation for its performance (**Supplementary Figure 6; Supplementary Table 6**). The novel-50 dataset was partitioned into two subsets using a conjunctive BLAST filter at ≥80% identity/ ≥80% coverage^27^. There were 42 targets that passed this filter, with the 8 targets (SARS-CoV-2 spike-antibody complexes) being the BLAST-flagged targets. *ProteinDock* reached 100% success rate on the full set, the pair-novel subset (Wilson 95% CI 91.6-100%), and the BLAST-flagged subset. This performance held at 70% and 90% identity thresholds as well, further indicating the result does not depend on the choice of identity threshold. We tested whether further reliability could be achieved within ref2015’s scoring space itself (**Supplementary Table 7**). Across the four model classes, only one physics correction (an interface-size based proxy) gave a positive but non-significant lift (**Supplementary Figure 7**). The refit of ref2015 weights had significant regression in performance (ΔDockQ = −0.060, p < 10^-4^) and conventional ML rescoring gave no meaningful gains. A set-transformer ensemble marginally improved decoy ranking (Spearman Δρ = +0.059) and top-1 selection (success rate 88.0% to 89.4%). Together, these results place a ceiling near 89% success rate, suggesting further improvement may require different information modalities beyond plain energy term modifications.

## Discussion

*ProteinDock* Mode 1 can be used in combination with Rosetta to improve reliability of protein-protein docking accuracy. Although improvement over standalone Rosetta was significant across sets, the reliability of *ProteinDock*-layered Rosetta was greater on antibody-antigen datasets than the comprehensive dataset, DB 5.5. This could be attributed to the distinct interface chemistry of antibody-antigen complexes, as well as similar performance trends of baseline Rosetta (as those output structures are what is refined by *ProteinDock*) (**Supplementary Figure 8**). Analysis of what interaction types fail on Rosetta and *ProteinDock* could motivate the development of *ProteinDock* variants that are optimized for specific protein-protein interaction types. Future studies can evaluate whether *ProteinDock* can be layered onto additional physics-based tools, such as HADDOCK, to improve the reliability of protein-protein docking.

Evaluation on an antibody-antigen set confirmed that the standard configuration of *ProteinDock* does not improve prediction reliability of AlphaFold3 for this interaction type. During an initial effort to optimize *ProteinDock* for this task, we found that a truncated version of *ProteinDock*, consisting only of the rerank step, can be used to evaluate outputs from multiple models and select the strongest one, ultimately improving reliability over a single model alone. This rerank step scores each candidate using *InterfaceAnalyzerMover* with the modified ref2015 function. When we compared this modified scoring against the standard *InterfaceAnalyzerMover* to rerank the outputs of AlphaFold3 and Boltz-2, the two performed with comparable reliability. This indicates the modification provides no additional advantage for this specific reranking. In this configuration, *ProteinDock* was tested on an antigen-antibody set. Future work should be aimed at continuing to optimize *ProteinDock* to improve prediction reliability for sequence-based predictors like AlphaFold3, building on the promising results of the truncated reranker. Comparing the truncated *ProteinDock* against standard *InterfaceAnalyzerMover* on more comprehensive datasets, or on datasets of alternative protein-protein interaction types, would help to establish the performance of this overall approach.

## Conclusion

We have developed *ProteinDock*, a Rosetta-based tool that uses a modified ref2015 score function to improve the protein-protein prediction reliability of leading programs. When docking from structure inputs with Rosetta, *ProteinDock* Mode 1 can be layered onto processes and rescore output structures. This setup generated 32.8% improvement over standalone Rosetta when evaluated on the Docking Benchmark set 5.5. Using Mode 1 of *ProteinDock* to process and rescore structural predictions from AlphaFold3 did not improve reliability when evaluated on the Novel-50 antibody-antigen set. However, a truncated *ProteinDock* Mode 2 setup, consisting solely of a rerank step, was used to select the best prediction between AlphaFold3 and Boltz-2, yielding a 6% improvement over standalone single-seed AlphaFold3. For ease of use and widespread accessibility, a graphical user interface has been created for layering *ProteinDock* and is accessible at https://github.com/Kimmel-lab/proteindock and https://www.proteindock.com.

## AUTHOR INFORMATION

### Corresponding Author

Blaise R. Kimmel, Ph.D. 151 W. Woodruff Avenue Columbus, OH 43210

### Author Contributions

The manuscript was written with contributions from all authors. All authors have approved the final version of the manuscript. B.R.K. supervised the project and acquired funding to support the research.

## Acknowledgements

This work was supported in part by The Ohio State University Center for Cancer Engineering, Curing Cancer Through Research in Engineering and Sciences, and the National Cancer Institute (NCI) through the R21 grant 1R21CA312456. B.R.K. acknowledges financial support from the Prostate Cancer Foundation Young Investigator Award. This work used the Ohio Supercomputer Center, which provides High Performance Computing resources and expertise to academic researchers across the State of Ohio. OSC is a member of the Ohio Technology Consortium, a division of the Ohio Department of Higher Education.

## References

(1) Chen, Y.; Clay, N.; Phan, N.; Lothrop, E.; Culkins, C.; Robinson, B.; Stubblefield, A.; Ferguson, A.; Kimmel, B. R. Molecular Matchmakers: Bioconjugation Techniques Enhance Prodrug Potency for Immunotherapy. Mol. Pharmaceutics 2025, 22 (1), 58–80. 10.1021/acs.molpharmaceut.4c00867.

(2) Bailey, J. S.; Spina, S. C.; Hu, A.; Phan, N.; Getman, R. B.; Kimmel, B. R. Artificial Intelligence in Chemical Engineering: Protein Design from First Principles to Structural Prediction. ACS Eng. Au 2026, acsengineeringau.5c00099. 10.1021/acsengineeringau.5c00099.

(3) De Chadarevian, S. John Kendrew and Myoglobin: Protein Structure Determination in the 1950s. Protein Science 2018, 27 (6), 1136–1143. 10.1002/pro.3417.

(4) Durham, J.; Zhang, J.; Humphreys, I. R.; Pei, J.; Cong, Q. Recent Advances in Predicting and Modeling Protein–Protein Interactions. Trends in Biochemical Sciences 2023, 48 (6), 527–538. 10.1016/j.tibs.2023.03.003.

(5) Leaver-Fay, A.; Tyka, M.; Lewis, S. M.; Lange, F.; Thompson, J.; Jacak, R.; Kaufmann, K. W.; Renfrew, P. D.; Smith, C. A.; Sheffler, W.; Davis, I. W.; Cooper, S.; Treuille, A.; Mandell, D. J.; Richter, F.; Ban, Y.-E. A.; Fleishman, S. J.; Corn, E.; Kim, D. E.; Lyskov, S.; Berrondo, M.; Mentzer, S.; Popovic, Z.; Havranek, J. J.; Karanicolas, J.; Das, R.; Meiler, J.; Kortemme, T.; Gray, J. J.; Kuhlman, B.; Baker, D.; Bradley, P. ROSETTA3: An Object-Oriented Software Suite for the Simulation and Design of Macromolecules.

(6) Raveh, B.; London, N.; Zimmerman, L.; Schueler-Furman, O. Rosetta FlexPepDock Ab-Initio: Simultaneous Folding, Docking and Refinement of Peptides onto Their Receptors. PLoS ONE 2011, 6 (4), e18934. 10.1371/journal.pone.0018934.

(7) Abramson, J.; Adler, J.; Dunger, J.; Evans, R.; Green, T.; Pritzel, A.; Ronneberger, O.; Willmore, L.; Ballard, A. J.; Bambrick, J.; Bodenstein, S. W.; Evans, D. A.; Hung, C.-C.; O’Neill, M.; Reiman, D.; Tunyasuvunakool, K.; Wu, Z.; Žemgulytė, A.; Arvaniti, E.; Beattie, C.; Bertolli, O.; Bridgland, A.; Cherepanov, A.; Congreve, M.; Cowen-Rivers, A. I.; Cowie, A.; Figurnov, M.; Fuchs, F. B.; Gladman, H.; Jain, R.; Khan, Y. A.; Low, C. M. R.; Perlin, K.; Potapenko, A.; Savy, P.; Singh, S.; Stecula, A.; Thillaisundaram, A.; Tong, C.; Yakneen, S.; Zhong, E. D.; Zielinski, M.; Žídek, A.; Bapst, V.; Kohli, P.; Jaderberg, M.; Hassabis, D.; Jumper, J. M. Accurate Structure Prediction of Biomolecular Interactions with AlphaFold 3. Nature 2024, 630 (8016), 493–500. 10.1038/s41586-024-07487-w.

(8) Hitawala, F. N.; Gray, J. J. What Does AlphaFold3 Learn about Antibody and Nanobody Docking, and What Remains Unsolved? mAbs 2025, 17 (1), 2545601. 10.1080/19420862.2025.2545601.

(9) Harmalkar, A.; Lyskov, S.; Gray, J. J. Reliable Protein–Protein Docking with AlphaFold, Rosetta, and Replica Exchange. eLife 2025, 13, RP94029. 10.7554/eLife.94029.

(10) Vreven, T.; Moal, I. H.; Vangone, A.; Pierce, B. G.; Kastritis, P. L.; Torchala, M.; Chaleil, R.; Jiménez-García, B.; Bates, P. A.; Fernandez-Recio, J.; Bonvin, A. M. J. J.; Weng, Z. Updates to the Integrated Protein–Protein Interaction Benchmarks: Docking Benchmark Version 5 and Affinity Benchmark Version 2. Journal of Molecular Biology 2015, 427 (19), 3031–3041. 10.1016/j.jmb.2015.07.016.

(11) Dunbar, J.; Krawczyk, K.; Leem, J.; Baker, T.; Fuchs, A.; Georges, G.; Shi, J.; Deane, C. M. SAbDab: The Structural Antibody Database. Nucl. Acids Res. 2014, 42 (D1), D1140–D1146. 10.1093/nar/gkt1043.

(12) Burley, S. K.; Berman, H. M.; Kleywegt, G. J.; Markley, J. L.; Nakamura, H.; Velankar, S. Protein Data Bank (PDB): The Single Global Macromolecular Structure Archive. In Protein Crystallography; Wlodawer, A., Dauter, Z., Jaskolski, M., Eds.; Methods in Molecular Biology; Springer New York: New York, NY, 2017; Vol. 1607, pp 627–641. 10.1007/978-1-4939-7000-1_26.

(13) Raghava, G.; Barton, G. J. Quantification of the Variation in Percentage Identity for Protein Sequence Alignments. BMC Bioinformatics 2006, 7 (1), 415. 10.1186/1471-2105-7-415.

(14) Schmitz, S.; Schmitz, E. A.; Crowe, J. E.; Meiler, J. The Human Antibody Sequence Space and Structural Design of the V, J Regions, and CDRH3 with Rosetta. mAbs 2022, 14 (1), 2068212. 10.1080/19420862.2022.2068212.

(15) Park, H.; Bradley, P.; Greisen, P.; Liu, Y.; Mulligan, V. K.; Kim, D. E.; Baker, D.; DiMaio, F. Simultaneous Optimization of Biomolecular Energy Functions on Features from Small Molecules and Macromolecules. J. Chem. Theory Comput. 2016, 12 (12), 6201–6212. 10.1021/acs.jctc.6b00819.

(16) Alford, R. F.; Leaver-Fay, A.; Jeliazkov, J. R.; O’Meara, M. J.; DiMaio, F. P.; Park, H.; Shapovalov, M. V.; Renfrew, P. D.; Mulligan, V. K.; Kappel, K.; Labonte, J. W.; Pacella, M. S.; Bonneau, R.; Bradley, P.; Dunbrack, R. L.; Das, R.; Baker, D.; Kuhlman, B.; Kortemme, T.; Gray, J. J. The Rosetta All-Atom Energy Function for Macromolecular Modeling and Design. J. Chem. Theory Comput. 2017, 13 (6), 3031–3048. 10.1021/acs.jctc.7b00125.

(17) Mohan, S.; Sinha, N.; Smith-Gill, S. J. Modeling the Binding Sites of Anti-Hen Egg White Lysozyme Antibodies HyHEL-8 and HyHEL-26: An Insight into the Molecular Basis of Antibody Cross-Reactivity and Specificity. Biophysical Journal 2003, 85 (5), 3221–3236. 10.1016/S0006-3495(03)74740-7.

(18) Boltz-2: Towards Accurate and Efficient Binding Affinity Prediction.

(19) Gray, J. J.; Moughon, S.; Wang, C.; Schueler-Furman, O.; Kuhlman, B.; Rohl, C. A.; Baker, D. Protein–Protein Docking with Simultaneous Optimization of Rigid-Body Displacement and Side-Chain Conformations. Journal of Molecular Biology 2003, 331 (1), 281–299. 10.1016/S0022-2836(03)00670-3.

(20) Bennett, N. R.; Watson, J. L.; Ragotte, R. J.; Borst, A. J.; See, D. L.; Weidle, C.; Biswas, R.; Yu, Y.; Shrock, E. L.; Ault, R.; Leung, P. J. Y.; Huang, B.; Goreshnik, I.; Tam, J.; Carr, K. D.; Singer, B.; Criswell, C.; Wicky, B. I. M.; Vafeados, D.; Sanchez, M. G.; Kim, H. M.; Vázquez Torres, S.; Chan, S.; Sun, S. M.; Spear, T.; Sun, Y.; O’Reilly, K.; Maris, J. M.; Sgourakis, N. G.; Melnyk, R. A.; Liu, C. C.; Baker, D. Atomically Accurate de Novo Design of Antibodies with RFdiffusion. Bioengineering March 18, 2024. 10.1101/2024.03.14.585103.

(21) Mirabello, C.; Wallner, B. DockQ v2: Improved Automatic Quality Measure for Protein Multimers, Nucleic Acids, and Small Molecules. Bioinformatics 2024, 40 (10), btae586. 10.1093/bioinformatics/btae586.

(22) Lensink, M. F.; Raouraoua, N.; Brysbaert, G.; Velankar, S.; Wodak, S. J.; Bonvin, A. M. J. J. Biomolecular Interaction Prediction in the Pre- and Post-ALPHAFOLD Era: The 8th CAPRI Evaluation. Proteins 2025, prot.70018. 10.1002/prot.70018.

(23) Mirdita, M.; Schütze, K.; Moriwaki, Y.; Heo, L.; Ovchinnikov, S.; Steinegger, M. ColabFold: Making Protein Folding Accessible to All. Nat Methods 2022, 19 (6), 679–682. 10.1038/s41592-022-01488-1.

(24) Giulini, M.; Reys, V.; Teixeira, J. M. C.; Jiménez-García, B.; V. Honorato R.; Kravchenko, A.; Xu, X.; Versini, R.; Engel, A.; Verhoeven, S.; Bonvin, A. M. J. J. HADDOCK3: A Modular and Versatile Platform for Integrative Modeling of Biomolecular Complexes. J. Chem. Inf. Model. 2025, 65 (13), 7315–7324. 10.1021/acs.jcim.5c00969.

(25) Jr, P. C. S.; Jacquemin, M.; Saint-Remy, J.-M. R.; Stoddard, B. L.; Pratt, K. P. Structure of a Factor VIII C2 Domain–Immunoglobulin G4k Fab Complex: Identification of an Inhibitory Antibody Epitope on the Surface of Factor VIII.

(26) Dussupt, V.; Jensen, J. L.; Rosado, A. P.; Donofrio, M.; Pflugheber, J.; Mendez-Rivera, L.; Sankhala, R. S.; Chen, W.-H.; Slike, B. M.; Schmid, A.; Tran, U.; Metzger, L.; Peterson, C. E.; Pinto, A. K.; Vasan, S.; Collins, N. D.; Farmer, A.; Michael, N. L.; Joyce, M. G.; Brien, J. D.; Krebs, S. J. Targeting the Zika Virus Envelope Domains I and III as a Recombinant Vaccine Protects Mice from Lethal Challenge. npj Vaccines 2026, 11 (1), 118. 10.1038/s41541-026-01442-8.

(27) Madden, T. The BLAST Sequence Analysis Tool.

