## Supplementary material for "ProteinDock: A physics-informed layer to improve protein-protein docking reliability": SI

**Supplementary Information**

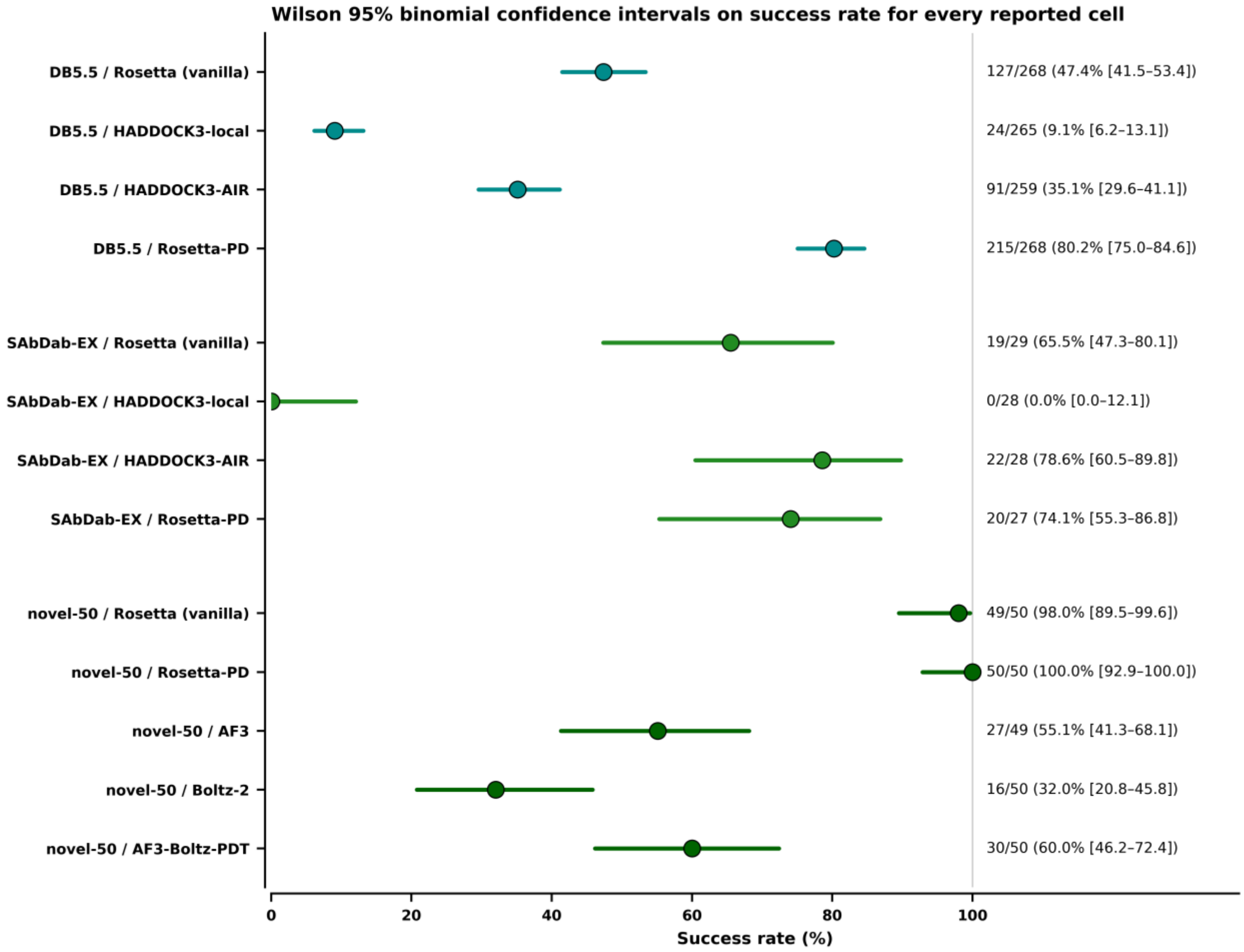


**Supplementary Figure 1.** Wilson 95% binomial confidence intervals on top-1 success rate for all methods and benchmark cells reported from the main text.

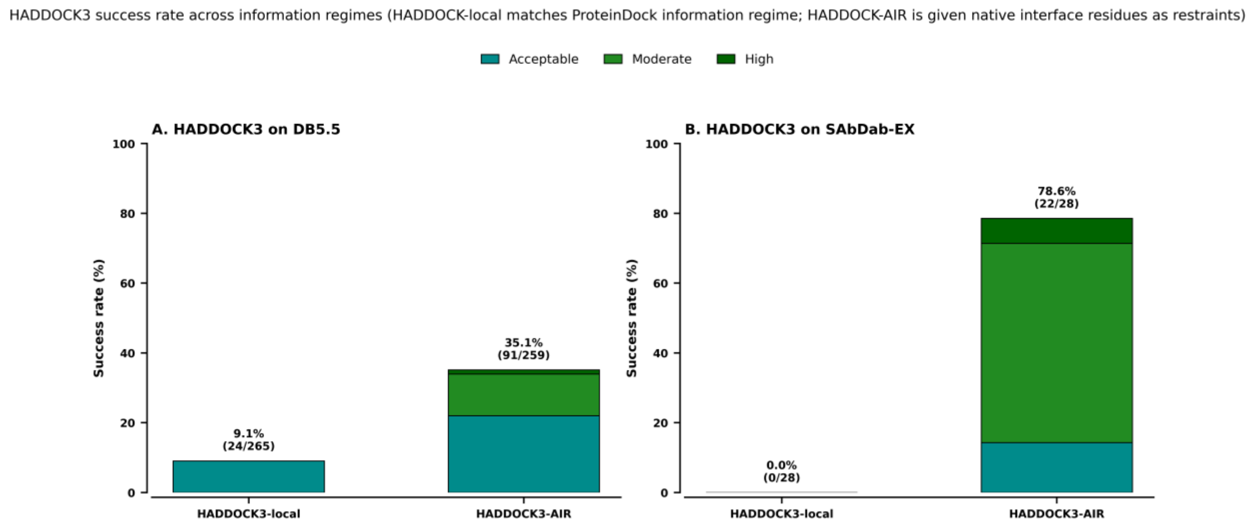


**Supplementary Figure 2.** HADDOCK3 success rate across HADDOCK3-local and HADDOCK-AIR for DB5.5 and SAbDab-EX.

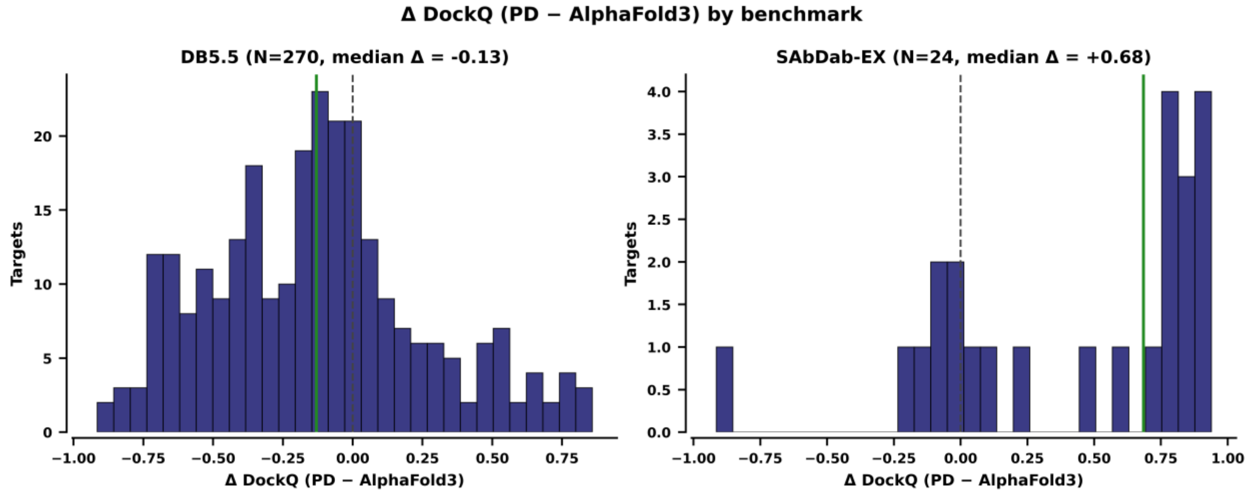


**Supplementary Figure 3.** Distribution of per-target ∆DockQ (ProteinDock-AlphaFold3) side by side with DB5.5 and SAbDab-EX.

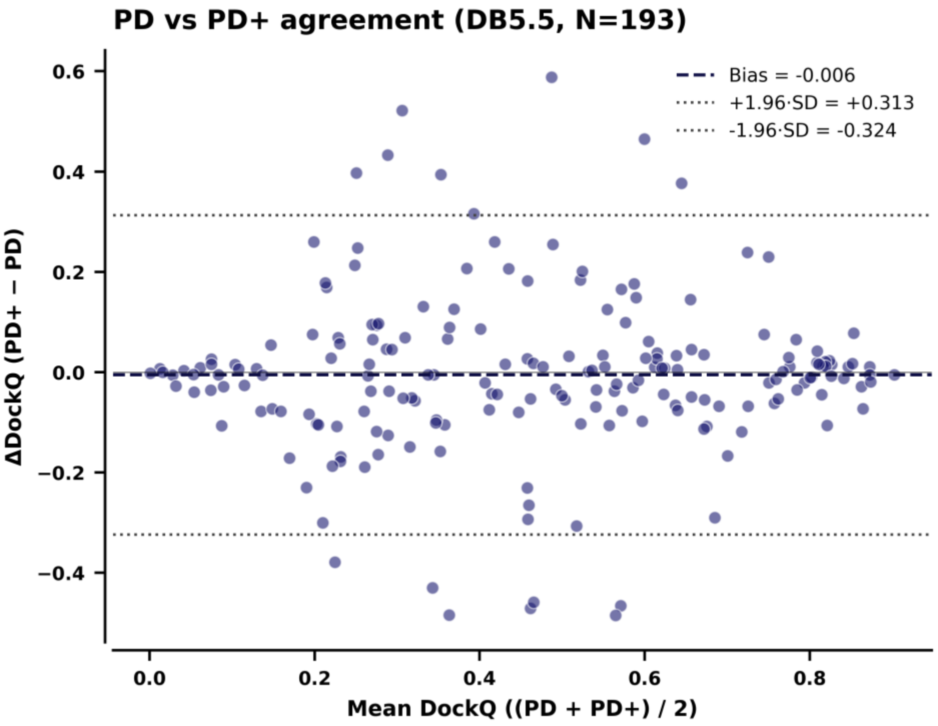


**Supplementary Figure 4.** Bland-Altman agreement between ProteinDock (PD) and PD+ per-target DockQ on paired DB5.5 targets, displaying mean bias and standard deviation limits.

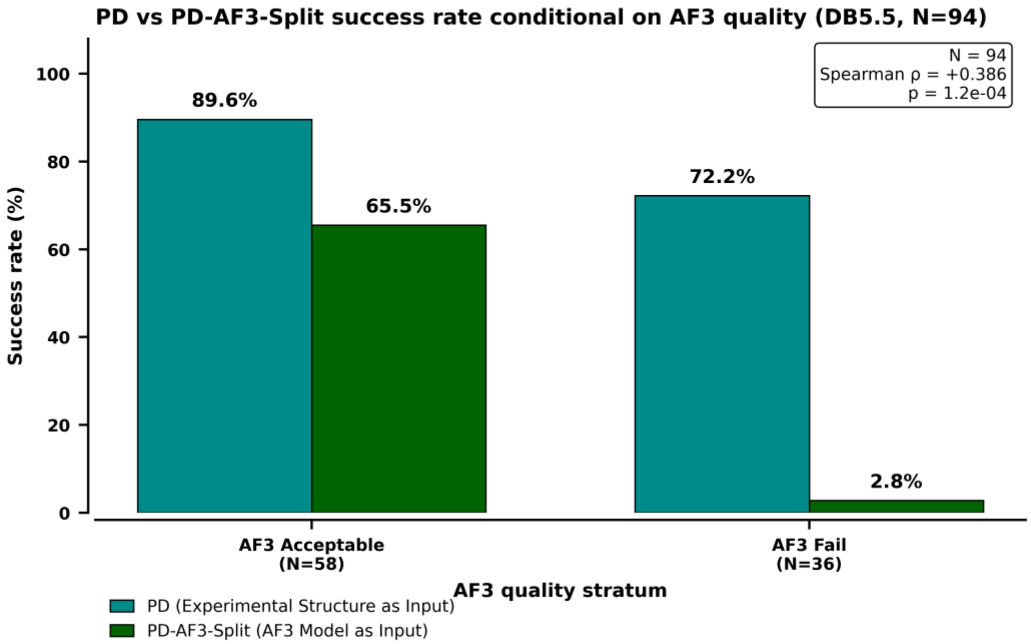


**Supplementary Figure 5.** ProteinDock success rate conditional on AlphaFold3 pose quality (DB5.5, N=94), comparing PD with PD-AF3-Split (AlphaFold3 model as input).

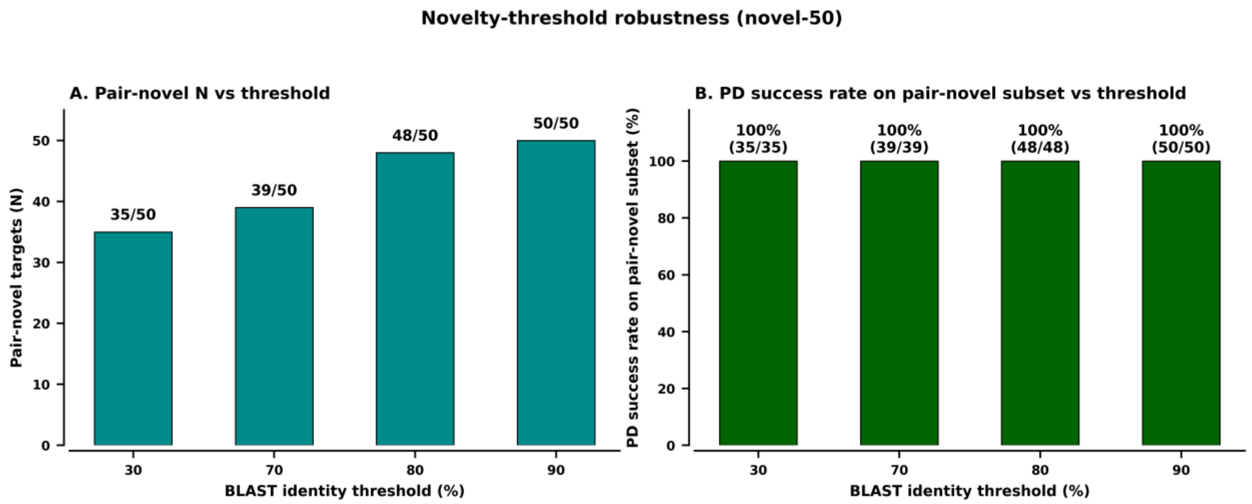


**Supplementary Figure 6.** Novelty-threshold robustness on novel-50. (A) Number of pair-novel targets vs BLAST chain-identity threshold. (B) ProteinDock top-1 success rate on the pair-novel subset at each threshold.

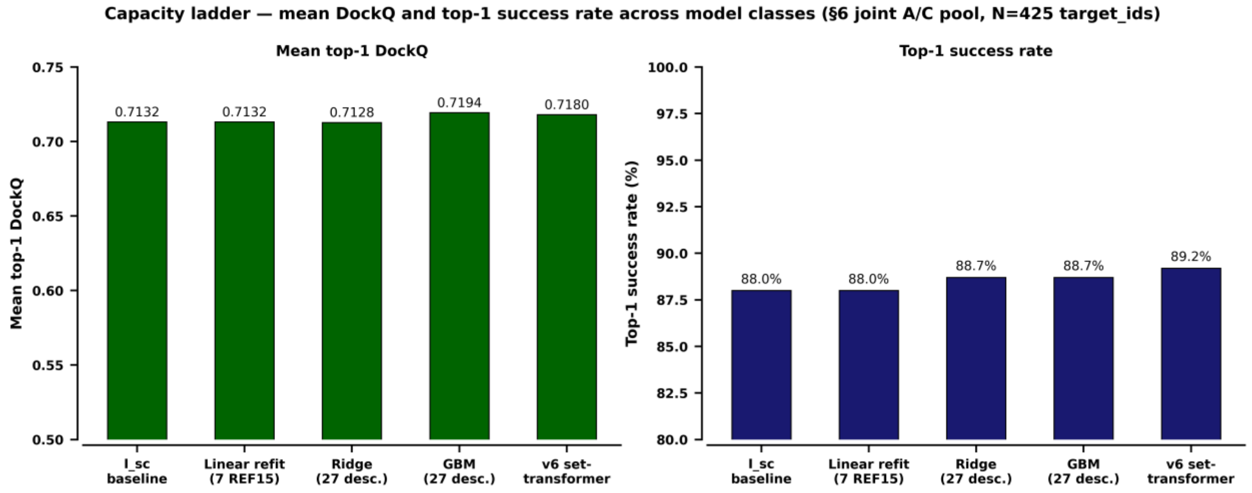


**Supplementary Figure 7.** Model-capacity ladder across different designs to assess ref2015-scoring capacity.

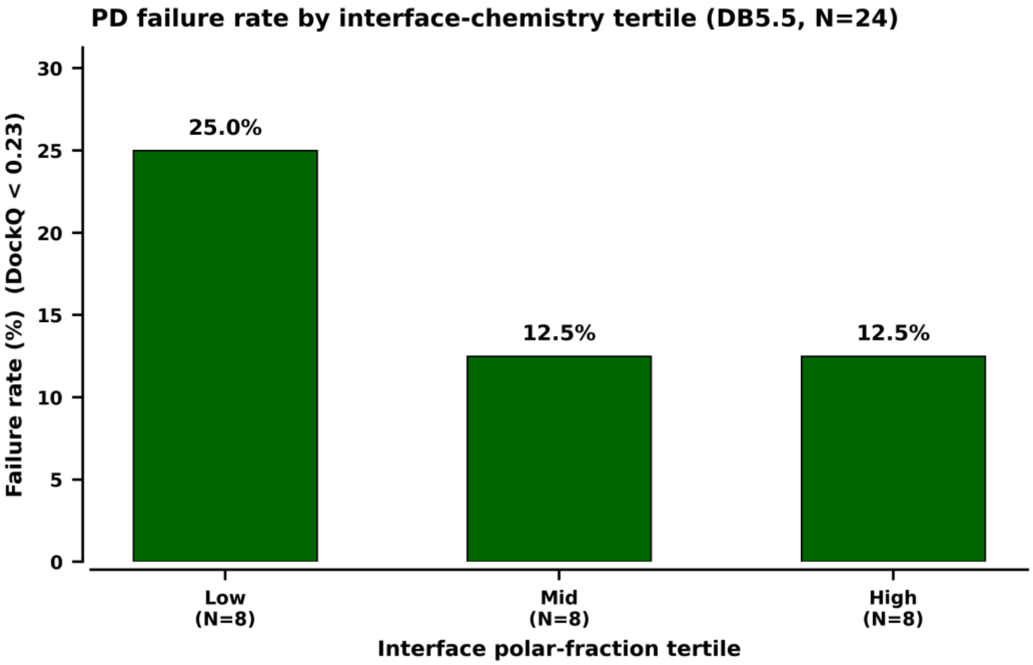


**Supplementary Figure 8.** PD failure rate (top-1 DockQ < 0.23) on DB5.5 separated into equal-sized tertiles of mean interface polar fractions.

**Supplementary Table 1.** Dataset curation.

| | DB5.5 | SAbDab-EX (SAbDab-Expanded) | novel-50 |
|---|---|---|---|
| Source | (Docking Benchmark; Vreven et al., 2015) | 20 complexes from SAbDab (Structural Antibody Database; Dunbar et. Al, 2014) + 9 complexes recovered from DB5.5 | PDB |
| Version/snapshot | 2015 release | October 2024 | Depositions 2021-10-01 to 2026-04-15 |
| N | 271 | 29 | 50 |
| Inclusion | Unbound structures available for both partners | Heterodimeric antibody-antigen pairs | Heterodimeric antibody-antigen pairs with paired H and L chains |
| Exclusion | —— | Complexes containing nucleic acid components | Antibody-antibody pairs; engineered crystal contacts |
| Resolution | No explicit threshold set | ≤ 2.8 Å | ≤ 3.0 Å (relaxed from ≤ 2.8 Å to reach N=50; 39 of 50 targets meet ≤ 2.8 Å) |
| Data Range | 1995-2015 | —— | Post-AlphaFold3 cutoff (2021-09-30) |
| Overlap | 27 of DB5.5's antibody-antigen entries appear in SAbDab-EX | 27 of 29 also in DB5.5; 1EZV and 1G9M are SAbDab only | No overlap with SAbDab-EX |

Pair-novelty was calculated with *BLAST+ 2.16.0* (blastp –evalue 1e-5) against 660,533 chain sequences from all 185,986 PDB entries from before 2021-09-30. A chain hit was kept if it reached at least 80% of the query length. A binding pair was counted as seen (and omitted from the pair-novel set) if at least one pre-cutoff PDB entry contained both a quality antigen hit and a qualifying antibody-chain hit in the same structure

**Supplementary Table 2.** Structure preparation by stage.

| | DockQ native | Mode 1 input | Mode 2 predictor pose |
|---|---|---|---|
| Waters/ ligands/ ions | Removed | Removed at chain preparation (fix_relabel_renumber.py strips water, ligands, and ions prior to FastRelax | n/a |
| Chains retained | Only protein chains listed in the benchmark index | Receptor + binder | As given by the predictor |
| Non-standard amino acids | Mapped to the canonical 20 | Removed (ACE/NME caps, hetatoms, modified residues stripped before combining; using only standard 20 amino acids) | n/a |
| Hydrogens | Not added prior to scoring | Stripped on preprocessing; FastRelax re-adds hydrogens under ref2015 default during pre-relax step | Not added |
| Chain relabelling | —— | Receptor: A; Binder: B . Multi-chain binders (antibody H+L) merged into a single relabeled binder | Receptor: A; Binder: B |
| Residue Renumbering | —— | Reset to sequential intergers per relabeled chain | Renumbered before scoring |
| Script | fix_relabel_renumber.py | fix_relabel_renumber.py | fix_relabel_renumber.py |

Receptor and binder chains were assigned per target with *Rosetta's* '-partners A_B' flag, with antibody H and L chains being specified as belonging to the same binder group with the purpose of maintaining CDR-framework geometry even through pre-relaxation. The system completes chain-labeling and scoring independently for each candidate per run to prevent cache bias in the final output.

**Supplementary Table 3.** Target counts by analysis.

| Analysis | Dataset(s) | N targets | Subset | Why this subset | N decoys |
|---|---|---|---|---|---|
| Mode 1 primary | DB5.5 | 271 | All | —— | 9722 given |
| Mode 1 primary | SAbDab-EX | 29 | All | —— | nstruct=50 requested; 1,125 delivered |
| Mode 1 primary | novel-50 | 50 | All | —— | nstruct=50 requested; 1,756 delivered |
| Pair-novelty | novel-50 | 42 | Pair-novel | Absent from pre-cutoff PDB analyzed by BLAST | —— |
| fa_elec sensitivity sweep | novel-50 | 42 | Validation subset | Test best value for fa_elec | —— |
| AF3-refinement runs | novel-50 | 42 | AF3-scorable | AF3 produced a scorable pose | —— |
| Reliability layer | novel-50 | 50 | All | Failures remain in the denominator | 25/target (AF3 pool) |
| Capture radius | novel-50 | 15 | 15 targets sampled uniformly across novel-50 | —— | 50 per cell x 60 cells |
| Capacity sweep pool | novel-50 + SAbDab + DB5.5 | 75 unique antibody-antigen complex PDBs | novel-50 (50) + SAbDab (18) + DB5.5 recovered (7) | —— | 13,240 across 425 groups |

**Supplementary Table 4A.** Variant specification (full), mean variants.

| Variant | Mode | Input Source | Pre-relax | Sampling | Rerank | Selection | Role |
|---|---|---|---|---|---|---|---|
| PD | 1 | Unbound (DB5.5, SAbDab-3), bound-frame monomers (novel-50) | FastRelax, ref2015 default weights, no constraints | nstruct=50, local docking @ fa_elec x 1.5 | —— | Top-1 by initial I_sc | Baseline docking method |
| AF3-PD | 2 | AF3's 25 seed x sample poses | —— | None | —— | argmin dG_interface | Single-predictor rescoring |
| AF3-Boltz-PDT | 2 | AF3 top-1 + Boltz-2 top-1 (single-seed protocol) | —— | None | —— | argmin with dG_interface, with fallbacks* | Cross-predictor arbitration |

*AF3-Boltz-PDT fallback logic: (1) If AF3 and Boltz-2 produced top-1 scorable top-1 poses, take the pose with the lower dG_interface value. (2) If AF3 produces a pose that Rosetta is unable to score, keep AlphaFold3's top-1 pose regardless. (3) If AF3 did not produce any output due to inference crashes, fall back to Boltz-2's top-1 pose.

**Supplementary Table 4B.** Variant specification (full), ablation variants

| Variant | Mode | Input Source | Pre-relax | Sampling | Rerank | Selection | Result (success rate) |
|---|---|---|---|---|---|---|---|
| PD+ | 1 | Unbound (DB5.5, SAbDab-3), bound-frame monomers (novel-50) | ✓ | nstruct=50 | Top-5 by initial I_sc, re-relaxed under same weights | Top-1 by post-relax I_sc | 50/50 (100% acc); High 82% |
| PD-AF3-Split | 1 | AF3 pose (top-1 by ranking_Score) | ✓ | nstruct=50 | —— | Top-1 by initial I_sc | 15/50 (30%) |
| PD-AF3-Split+ | 1 | AF3 pose (top-1 by ranking_Score) | ✓ | nstruct=50 | Top-5 FastRelax rerank | Top-1 by post-relax I_sc | 12/50 (24%) |
| PD-AF3-Polish | 1 | AF3 pose (top-1 by ranking_Score) | Coordinate-constrained FastRelax only | None | —— | —— | 13/50 (26%) |
| PD-AF3-Refine | 1 | AF3 pose (top-1 by ranking_Score) | —— | Tight rigid-bdoy refine, dock_pert 0.5 1.5, nstruct=25 | —— | Top-1 by I_sc | 0/50 (0%) |
| PD-AF3-Cluster | 1 | AF3 pose (top-1 by ranking_Score) | —— | Cluster into groups, refine each three ways (FastRelax polish, rigid-body refine, and re-docking), rerank, relax top 5 | Multi-metric | Top-1 by post-relax I_sc | 11/50 (22%) |
| PD-AF3-Dock | 1 | AF3 pose (top-1 by ranking_Score) | —— | Full-perturbation local docking | Multi-metric | Top-1 by post-relax I_sc | 0/0 (Did not run due to killing AF3 poses) |

**Supplementary Table 5A.** Baseline method specifications: AlphaFold3 (Abramson *et al.*, 2024)

| Parameter | Value |
|---|---|
| Release | Public release (Abramson et al., 2024), 2024-11-11 model weights |
| Mode | Protein-only |
| Inputs | Receptors and binder amino-acid sequence; multi-chain antibodies supplied as separate H and L chains |
| MSA | Generated internally from AlphaFold3's bundled database |
| Template filtering | Filtered to depositions on or before 2021-09-30 (matching training cutoff; verified per-target that retrieved pre-cutoff chain templates did not encode binding geometry |
| Seed/samples (DB5.5, SAbDab-EX) | 1 seed x 5 diffusion sanmples; highest 'ranking_score' sample taken as prediction |
| Seeds/samples (novel-50) | 5 seeds x 5 diffusion samples (25 candidates/target); highest 'ranking_score' sample across the pool taken as prediction |
| Hardware | 1x NVIDIA A100 (40 GB), Ohio Supercomputer Center (Pitzer/Cardinal) |
| Runtime | ~0.5 GPU-hours/target (DB5.5, SAbDab-EX); ~2.5 GPU-hours/target (novel-50, 5 seeds) |

**Supplementary Table 5B.** Baseline method specifications: Boltz-2 (Passaro *et al.,* 2025)

| Parameter | Value |
|---|---|
| Version | V2.2.1 (boltz CLI, pyPI) |
| Invocation | boltz predict |
| Seeds | 1 random seed per target |
| Inputs | Receptor and binder amino-acid sequences, YAML manifest |
| MSA | Use_msa_server: true<br><br>routed to ColabFold MMseqs2 server (Mirdita et al., 2022; uniref30 + colabfold-envdb-202108) |
| Sampling | --diffusion_samples 5, --recycling_steps 3 |
| Selection | Top-ranked (model_0) sample taken as per-target prediction |
| Post-processing | Predictions relabeled to chain A (receptor)/ chain B (binder) for DockQ comparison |
| Hardware | 1x NVIDIA A100 (40 GB), Ohio Supercomputer Center (Pitzer/Cardinal) |
| Runtime | ~0.1 GPU-hours/target |

**Supplementary Table 5C.** Baseline method specifications: HADDOCK3 (Giulini *et al.,* 2025)

| Parameter | HADDOCK-local | HADDOCK-AIR |
|---|---|---|
| Restraints | Chain coordinates only, no AIRs | Native interface residues as ambiguous interface residues (AIRs), extracted from the bound complex |
| Interface information | Matched to ProteinDock | Maximum (native contacts) |
| w_desolv | 0.0 | Default HADDOCK3 weights |
| Sampling | 100 rigid-body with cmrest →<br>40 flex →<br>Top-1 by HADDOCK score | 500 rigid-body with AIRs →<br>80 flex →<br>Top-1 by HADDOCK score |

Version: HADDOCKv3 v2026.5.0, run on DB5.5. Standard HADDOCK protocol stages were used throughout. Top-1 selection by HADDOCK score.

**Supplementary Table 5D.** Baseline method specifications: vanilla Rosetta (Leaver-Fay *et al.,* 2011)

| Parameter | Value |
|---|---|
| Definition of "vanilla" | No pre-relax step, no fa_elec reweight |
| Inputs | Identical to ProteinDock Mode 1 |
| Preprocessing | Nonstandard residues stripped before combining (ACE/NME caps, hetatoms, waters, ligands, modified residues) |
| Binary | rosetta_scripts.static.linuxgccrelease |
| Protocol | DockingProtocol (Gray et al., 2003) |
| Flags | -dock_pert 3 8, -partners A_B, -score:weights ref2015.wts, -nstruct 50 per target |
| Selection | Top-1 by Rosetta interface score (I_sc) from docking scorefile |
| Hardware | Intel Xeon Gold 6148 CPU, 4 cores/SLURM array task, Ohio Supercomputer Center (Pitzer/Cardinal) |
| Runtime | ~2 CPU-hours/target |

**Supplementary Table 6.** Data issues and corrections.

| Issue | Effect | Cause | Fix | Analyses Affected |
|---|---|---|---|---|
| 1EZV | Wrong antigen-chain assignment during scoring | SAbDab-EX processing | Corrected before evaluation | SAbDab-EX scoring pipeline |
| 1BVK | Empty antibody.pdb, no scorable decoys | Missing source file in original SAbDab curation | Rebuilt complex | Capacity-sweep pool; SAbDab-EX; Mode 1 decoys |
| Chain-merge/ numbering | DockQ returns iRMS = 999 even on correctly predicted structures | Numbering mismatch between merged binder and separately numbered native H/L | Binder residues renumbered to the union of native H and L indices, applied uniformly | All DockQ scoring |
| 7 misclassified DB5.5 complexes | Antibody-antigen complexes found in the general-PPI pool | Mis-annotation in DB5.5 | Recovery by hand and added to the capacity-sweep pool | Capacity-sweep pool |
| Non-converging structures | Rosetta docking exits with all decoys rejected | Steric clashes in input complex that FastRelax cannot resolve | Standard processing filter | Vanilla Rosetta DB5.5 (3 targets) |

**Supplementary Table 7.** Descriptor definitions.

| Name | Count | Members | Used in |
|---|---|---|---|
| Chemistry descriptors | 5 | Hydrophobic fraction, polar fraction, interface size (residues), charged residues, salt bridges | Antibody vs. non-antibody interface comparison, Mann-Whitney U (below) |
| Single-parameter physics candidates | 14 | Interface size (residues); interface size (heavy atoms); contact count (4.5 Å heavy-atom); contact count (5 Å heavy-atom); sum of Voronoi contact areas; number of hydrogen bonds; number of salt bridges; number of aromatic-aromatic contacts; number of hydrophobic-hydrophobic contacts; number of charged-charged contacts; per-CDR contact counts (H1, H2, H3 aggregated); per-CDR contact fractions (H1, H2, H3 aggregated); packing density (Voronoi cell fraction); clash count (< 3 Å heavy-atom) | Calibration-robustness (I_sc + α· X corrections) |
| ref2015 energy terms | 7 | fa_atr, fa_rep, fa_sol, fa_elec, hbond_sc, hbond_bb_sc, lk_ball_wtd | Linear refit; capacity sweep |
| Interface geometry features | 27 | Interaction counts within 4.5 Å (charged–charged, aromatic–aromatic, hydrophobic–hydrophobic, H-bond donor–acceptor, salt-bridge); Voronoi-style distance-weighted contact areas; per-CDR contact counts and fractions across all six CDR loops; clash counts; packing density | Capacity sweep, per-decoy |
| Target context features | 5 | Receptor size (residues); binder size (residues); interface class (Ab-Ag flag from index); antigen chain count; per-target decoy pool size | Capacity sweep, per-target |

**Supplementary Methods 1:** *Capture-radius characterization*

We perturbed 15 novel-50 targets from their native bound state across four different magnitudes: proximal (2 Å, 5°), near (5 Å, 10°), mid-range (8 Å, 15°), and far (12 Å, 20°), with 60 target-magnitude cells. Each perturbed input was run through the full ProteinDock protocol, and the final outputs were classified into the four CAPRI bins. We used one seed per cell due to the ability of Rosetta's DockingProtocol to sample randomly at each starting pose, already providing the variance an extra seed would otherwise add.

**Supplementary Methods 2:** *Capacity sweep, full specifications*

The capacity sweep was completed on the pool described in **Supplementary Table 3** under 5-fold PDB-disjoint cross-validation (descriptors in **Supplementary Table 7**).

| Model Class | Free Parameters | Training |
|---|---|---|
| Single-parameter physics correction, I_sc + α· X | 1 | α tuned per fold, grid search |
| Linear ref2015 weight refit | 7 | Pairewise margin-rank loss |
| Conventional regressors (27 variants) | $10^2$-$10^4$ | sklearn defaults |
| Set-transformer ensemble, 15 seeds | $10^5$ | AdamW, cosine annealing, early stopping* |

*Set-transformer architecture: 2-layer encoder, 4 heads, d_model= 96, dropout= 0.25, sigmoid DockQ head. Trained with 1:1 MSE + LambdaRank pairwise loss, AdamW (lr=1 x $10^{-3}$), cosine annealing over 120 epochs, gradient clipping at 1.0, early stopping (patience 15) on a 20% internal validation split.
